# Cellulase secretion by engineered *Pseudomonas putida* enables growth on cellulose oligomers

**DOI:** 10.1101/2025.06.19.659671

**Authors:** Madeline R. Smith, Kaylee Moffitt, William Holdsworth, Carlos H. Luna-Flores, Mansi Goyal, Alex Beliaev, Robert E. Speight, James B. Behrendorff

## Abstract

*Pseudomonas putida* is an attractive synthetic biology platform organism for chemical synthesis from low-grade feedstocks due to its high tolerance to chemical solvents and lignin-derived small molecules that are often inhibitory to other biotechnologically relevant microorganisms. However, there are few molecular tools available for engineering *P. putida* and other gram-negative bacteria to secrete non-native enzymes for extracellular feedstock depolymerisation. In this study *P. putida* was transformed to secrete cellulase enzymes and evaluated for growth on polymeric or oligomeric cellulose substrates. Active exo- and endocellulase enzymes were secreted into the culture supernatant, and a preferred set of twin-arginine translocase secretion signal peptides were identified. Extracellular cellulase activity was sufficient to support growth of *P. putida* using cellotriose or cellotetraose as the sole source of carbon and energy. This work supports progress towards consolidated bioprocessing of cellulosic materials using *P. putida*, and advances the state of engineered protein secretion in gram negative bacteria.

**Key Points:**

- Engineered *Pseudomonas putida* secreted cellulase enzymes into the culture medium
- Cellulase activity was sufficient to support growth on cellulose oligomers

## Introduction

Deployment of microbial fermentation for chemical synthesis at industrial scales depends on the ability to utilize low-cost feedstocks of varying composition and quality, such as low-value lignocellulosic crop residues or other macropolymeric materials. These resources require either pre-treatments to yield fermentable monomers, or consolidated bioprocessing by organisms that secrete enzymes for extracellular depolymerisation of high molecular weight material as well as converting the released monomers into valuable products.

Consolidated bioprocessing of cellulosic feedstocks explores the idea of combining cellulose hydrolysis and fermentation in a single step, where the fermentation organism also secretes cellulolytic enzymes to convert cellulose to glucose. Optimised consolidated bioprocessing can reduce the complexity and cost of biochemical manufacturing (Dempfle et al. 2021), but most bacteria and yeasts used in precision fermentation do not naturally secrete cellulases and few tolerate the inhibitory compounds typically present in crude preparations of cellulose from lignocellulosic biomass (den Haan et al. 2015). *Pseudomonas putida* is an emerging synthetic biology platform organism for fermentative biochemical synthesis (Weimer et al. 2020), and it is an attractive candidate for consolidated bioprocessing because it is highly tolerant to lignin-derived aromatic compounds and organic acids present in lignocellulosic biomass hydrolysates (Borchert et al. 2023). However, *P. putida* does not natively secrete cellulolytic enzymes and cannot grow on cellulose. In this study we aimed to establish secretion of cellulase enzymes by *P. putida* and evaluate the potential for developing consolidated bioprocessing with this organism.

Engineered secretion of enzymes in gram-negative bacteria (including *P. putida*) is relatively unexplored due to the complexity presented by gram-negative cell wall anatomy. In gram-negative bacteria, the general secretory (Sec) and twin arginine translocase (Tat) pathways recognise specific N-terminal signal sequences to chaperone proteins across the inner membrane into the periplasmic space (Costa et al. 2015). Extracellular secretion of heterologous proteins by *Escherichia coli* is largely dependent on the use of signal peptides that in most cases direct protein secretion to the periplasm, and extracellular protein release frequently depends on rupture of the outer membrane (Kleiner-Grote et al. 2018). While the molecular mechanisms of gram-negative secretion systems are well characterised, there are relatively few successful demonstrations of extracellular secretion in gram-negative bacteria and these are reliant on relatively few secretion signal peptides (Burdette et al. 2018). Functional validation of further secretion signal peptides in additional gram-negative hosts is needed.

Many proteins located on the extracellular surface of the outer membrane are transported via the autodisplay pathway, whereby a protein is first secreted to the periplasmic space via the Sec pathway and then a C-terminal β-barrel domain embeds into the outer membrane and chaperones the N-terminal functional domain to the extracellular surface, where it remains tethered. Autodisplay has been used to display cellulases on the surface of *P. putida* (Schulte et al. 2017), and to develop surface-displayed dockerins for *in vitro* assembly of extracellular cellulosome-like complexes (Dvořák et al. 2020).

In nature, surface-displayed cellulosomes have two primary benefits: assembly of a consortium of enzymes for hydrolysis of mixed polymeric feedstocks (Gilbert 2007), and adhesion of the microorganism to the insoluble polymeric substrate (Shoham et al. 1999). In a biorefinery context where defined microbial strains are grown in a closed fermentation, surface display of cellulolytic enzymes is unnecessary and may impose a sub-optimal upper limit on the concentration of cellulase enzyme. We engineered *P. putida* to secrete free cellulases by using secretion signal peptides associated with native extracellular proteins, but omitting hydrophobic anchoring domains that would otherwise link the cargo protein to the cell surface or vesicles.

## Results

### Designing cellulase secretion in Pseudomonas putida

Two Tat signal peptides and two Sec signal peptides associated with extracellular proteins were identified from the published literature. A characterised extracellular phosphatase, UxpB (Putker et al. 2013), is secreted via the Tat pathway, while the most abundant extracellular protein, outer membrane porin OprF (Choi et al. 2014), is secreted via the Sec pathway. Two additional extracellular proteins (one with a Sec secretion signal, Genbank locus WP_010955666.1, and one with a Tat signal, Genbank locus WP_010953415.1), were identified from a proteomic study of *P. putida* outer membrane vesicles (Salvachúa et al. 2020) (Online Resource 1, Supporting Table S1).

The CelA endocellulase and CelK exocellulase from *Acetivibrio thermocellus* (a gram-positive bacterium, formerly known as *Clostridium thermocellum*) (Tindall 2019) were selected as model cellulases. While thermostable, these enzymes have comparable activity at 30 °C to known mesophilic prokaryotic cellulases (Mingardon et al. 2011), of which few have been characterised. Cellulolytic fermentations with *A. thermocellus* (Shao et al. 2020) and assays of its cellulosome complex (Zhang and Lynd 2003) are typically conducted at or close to neutral pH, inidcating compatibility with *P. putida* culture conditions (initial medium pH of 7.0). Their native Sec pathway secretion signal peptide sequences of the *A. thermocellus* CelA and CelK cellulases were identified and replaced with the Sec or Tat pathway signal peptides from *P. putida* and expressed from a bicistronic expression plasmid under the control of the xylS/Pm inducible promoter system(Gawin et al. 2017) (plasmids pSEC_cellulase and pTAT_cellulase; Online Resource 1, Supporting Figure S1).

Expression constructs were designed with low translation initiation rates (TIRs; De Novo DNA Operon Calculator, http://www.denovodna.com) for each cellulase gene, using a TIR of approximately 10,000 units for the CelK exocellulase and 1000 units for the CelA endocellulase, on the basis that efficient release of glucose from a cellulose polymer requires more units of exocellulase activity than endocellulase (Setter-Lamed et al. 2017).

### Secretion of active cellulases from Pseudomonas putida

Extracellular cellulase activity was observed on solid media by plating transformed *P. putida* strains on M9 agar supplemented with carboxymethylcellulose (CMC; a non-fermentable water-soluble cellulose derivative) and 3-hydroxybenzoic acid (3-HB; inducer of the xylS/Pm promoter system). Agar plates were subsequently stained with Congo Red, which interacts strongly with intact β- (1-4)-glucopyranosides. Pale orange clearance zones indicating CMC hydrolysis were observed for *P. putida* KT2440 bearing the pTAT_cellulase and pSEC_cellulase plasmids, but not for strains bearing the negative control pSEVA231 plasmid (Figure 1). This observation was consistent when the *P. putida* S12 strain was transformed with the same plasmids (Online Resource 1, Supporting Figure S2).

**Figure 1.**
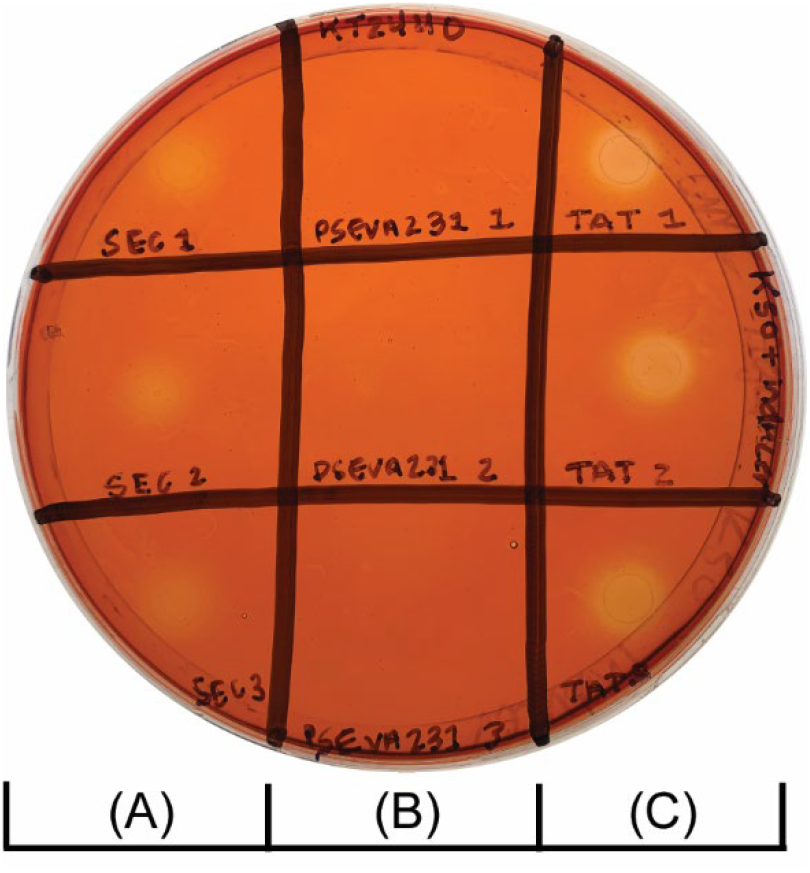
Carboxymethylcellulose hydrolysis on solid agar. Liquid cultures (5 μL) of *P. putida* KT2440 transformed with pSEC_cellulase (A), pSEVA231 (B), and pTAT_cellulase (C) plasmids were spotted on M9 agar containing carboxymethylcellulose (0.5% w/v) and kanamycin (50 µg/mL). Carboxymethylcellulose hydrolysis was visualised by staining with Congo Red. Three biological replicates for each strain are arranged in each marked column.

The presence of CelK and CelA cellulases in supernatants from pSEC_cellulase and pTAT_cellulase cultures was confirmed via targeted proteomics, where the CelK and CelA cellulases were identified as the presence of a peptide unique to each cellulase that was not observed in negative control strains bearing the pSEVA231 plasmid. The CelK cellulase was also qualitatively identified in the intracellular fraction, while intracellular CelA was not detected (Online Resource 1, Supporting Table S2).

Liquid culture supernatants were assessed for presence of functional cellulases using a panel of probe substrates. Hydrolysis of CMC was detected by reacting oligosaccharide reducing ends with *p*-hydroxybenzoic acid hydrazide (Lever 1972) (PAHBAH; Figure 2A). Cellulase activity in culture supernatants was also assayed with 4-methylumbelliferyl β-D-cellobioside (Chernoglazov et al. 1989) (4-MU cellobioside; Figure 2B) and *p*-nitrophenyl β-D-cellobioside (Deshpande et al. 1984) (*p*-NP cellobioside; Figure 2C), which yield fluorescent methylumbelliferone and coloured *p*-nitrophenol products, respectively, upon hydrolysis of the agluconic bond between the cellobiose group and the reporter molecule. Strains bearing plasmids designed for cellulase secretion catalysed significant hydrolysis of all three substrates, and in each case the rate of hydrolysis was greater with supernatants from pTAT_cellulase cultures. These observations were again consistent when repeated in the *P. putida* S12 strain (Online Resource 1, Supporting Figure S3).

**Figure 2.**
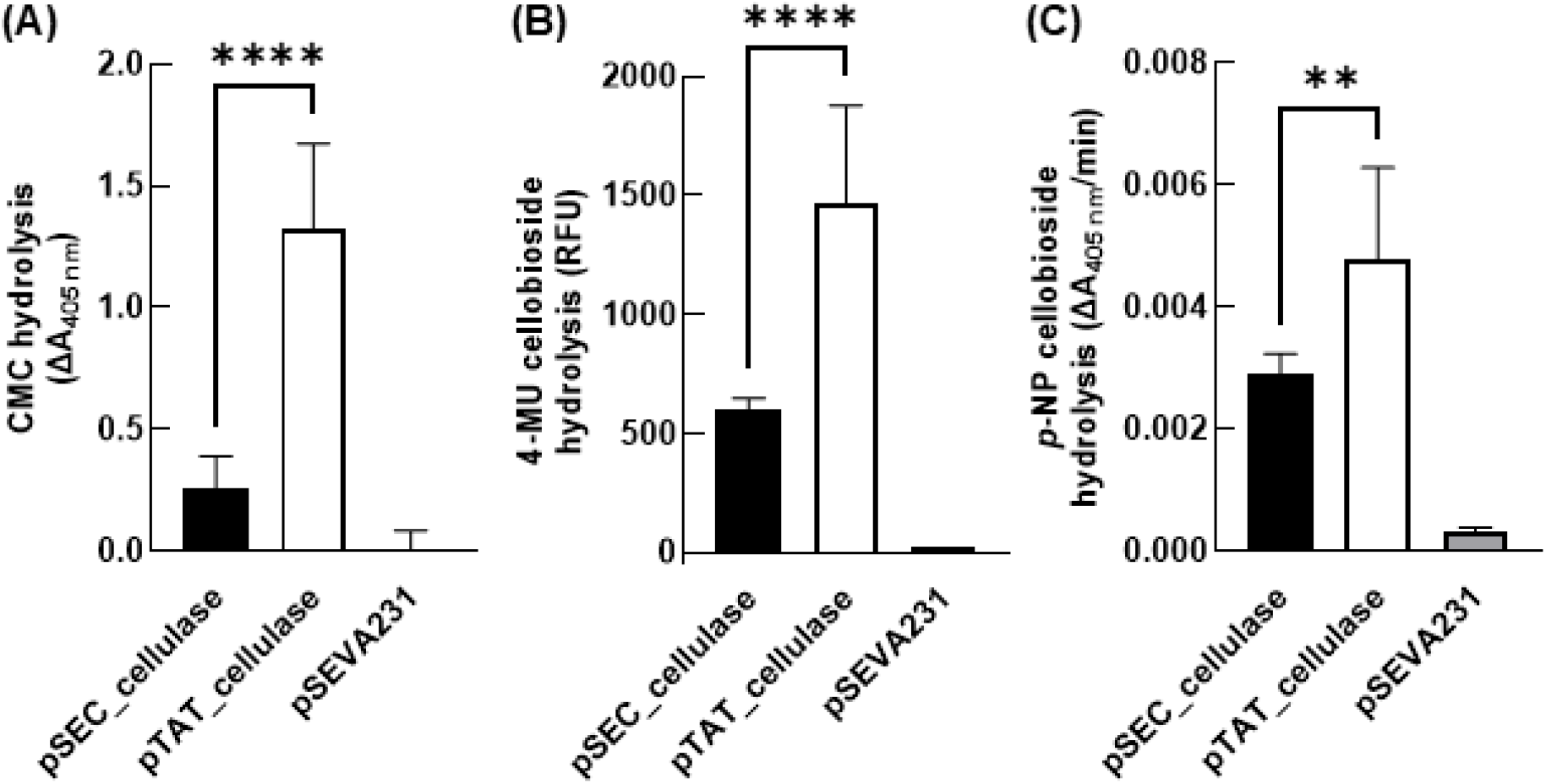
Cellulase activity in culture supernatants. *P. putida* KT2440 expression cultures were sedimented via centrifugation and the culture supernatants were incubated with a variety of probe substrates for detection of cellulase activity. (A) Carboxymethylcellulose (CMC) hydrolysis was assayed via reaction of reducing sugar ends with *p*-hydroxybenzoic acid hydrazide. (B) 4- methylumbelliferyl β-D-cellobioside (4-MU cellobioside) hydrolysis was assayed as fluorescence of liberated 4-methylumbelliferone. (C) *p*-nitrophenyl β-D-cellobioside (*p*-NP cellobioside) hydrolysis was assayed as change in absorbance from liberated *p*-nitrophenol, comparing rate of product formation in the initial linear phase of the reaction. All plots report the mean of n = 3 biological replicate cultures + standard deviation, unpaired Student’s *t*-test (**** = p < 0.0001, ** = p < 0.01).

Further plasmids were designed to independently investigate the functionality of the two identified TAT secretion signal peptides and the impact of the low TIR on the CelA endocellulase (Table 1). The CelK exocellulase was expressed using the same signal peptide (uxpB) and TIR as in the pTAT_cellulase plasmid. The CelA endocellulase was expressed with either the PP_2478 or the uxpB signal peptide, and at two different TIRs (∼1000 and ∼10,000). All CelA endocellulase variants produced similar CMC clearance zones to that observed for the pTAT_cellulase strain (Online Resource 1, Supporting Figure S4). A fainter clearance zone was observed when the CelK exocellulase was expressed alone.

**Table 1.** Key features of cellulase secretion plasmids used in this study.

| Plasmid name | CelK exocellulase | CelA endocellulase |
| --- | --- | --- |
| pSEVA231 | - | - |
| pTAT_cellulase | uxpB signal peptide, TIR 13356 | PP_2478 signal peptide, TIR 1346 |
| pSEC_cellulase | OprF signal peptide, TIR 10504 | PP_5130 signal peptide, TIR 1057 |
| pUxpB-celK_10k | uxpB signal peptide, TIR 13356 | - |
| p2478-celA_1k | - | PP_2478 signal peptide, TIR 1346 |
| pUxpB-celA_1k | - | uxpB signal peptide, TIR 1035 |
| p2478-celA_10k | - | PP_2478 signal peptide, TIR 10172 |
| pUxpB-celA_10k | - | uxpB signal peptide, TIR 9018 |

The CelA endocellulase demonstrated CMC hydrolysis with either the PP_2478 or uxpB secretion signal peptide (Supporting Figure S5), and CMC hydrolysis was roughly doubled when the celA endocellulase TIR was increased from ∼1000 to ∼10,000 units. Only the CelK exocellulase catalysed hydrolysis of 4-MU cellobioside and *p*-NP cellobioside (Supporting Figure S5).

### Pseudomonas putida growth on cellulose oligomers

A *P. putida* strain that constitutively expresses an intracellular β-glucosidase for metabolism of cellobiose, *P. putida* KT2440_P_*EM7*_-cbh, was prepared based on a previously-published design (Dvořák and de Lorenzo 2018) (Supporting Methods S1 and Supporting Figure S6). The pSEC_cellulase and pTAT_cellulase plasmids were separately used to transform *P. putida* KT2440_P_*EM7*_-cbh, and capacity for growth on glucose and cellotriose was assessed. Addition of 3-methybenzoic acid to induce gene expression caused a modest growth penalty across all strains when grown on in minimal media with glucose as the sole carbon source (Figure 3A). The greatest growth on cellotriose was observed in uninduced cultures of the pTAT_cellulase strain with co-expression of the intracellular cellobiohydrolase (pTAT_cellulase P_*EM7*_-cbh, Figure 3B, Supporting Figure S7). Marginal growth was observed pSEC_cellulase in the P_*EM7*_-cbh genetic background, whereas strains lacking the cellobiohydrolase were not viable when grown on cellotriose.

**Figure 3.**
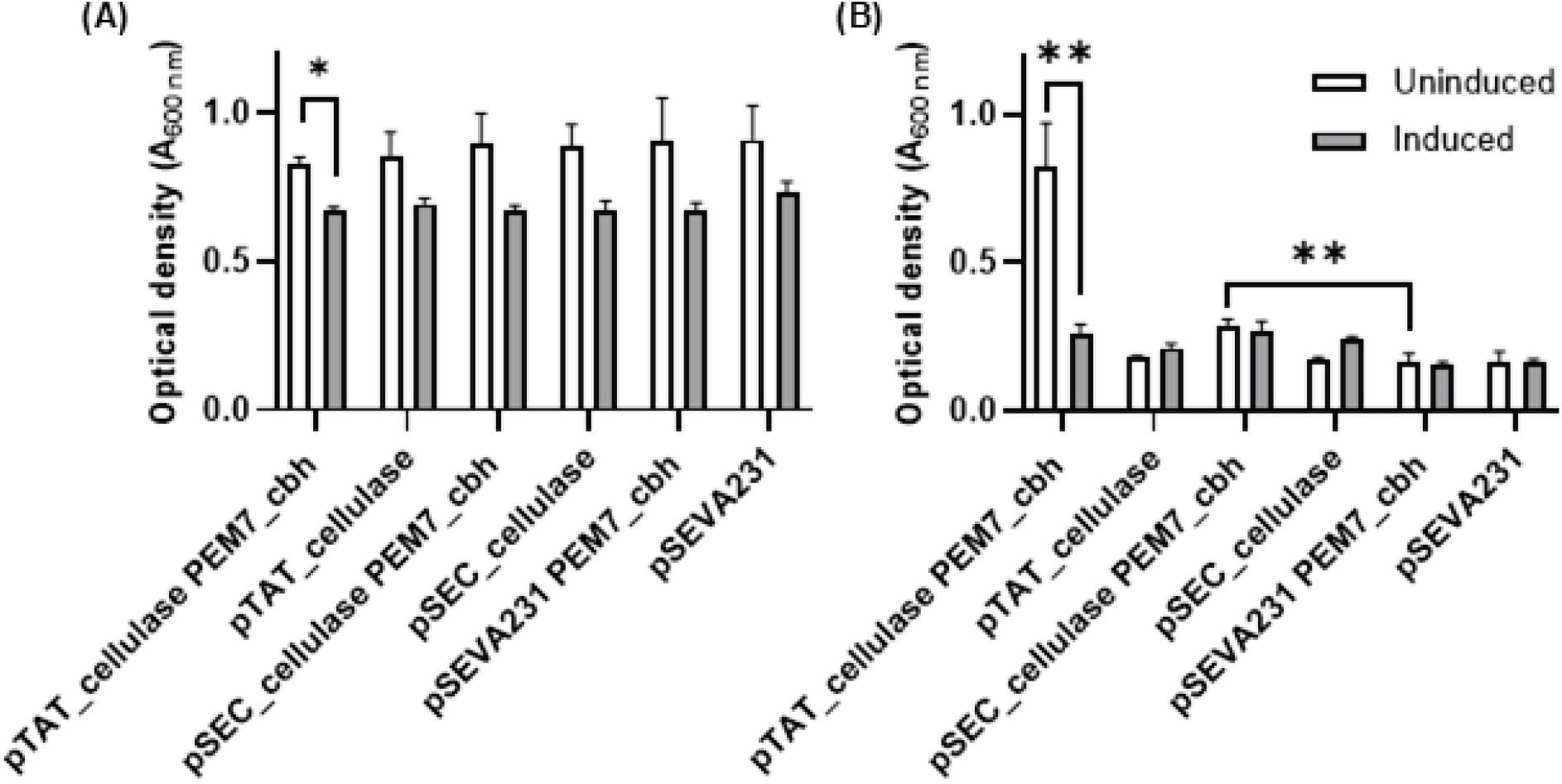
Growth with or without chemical induction of cellulase expression. *P. putida* cellulase expression strains were grown with (A) glucose or (B) cellotriose as the sole carbon source, either with chemical induction of gene expression (addition of 1 mM 3-methylbenzoic acid, grey bars) or without induction (white bars). The maximum optical density within a 120 h window was recorded (n = 3 biological replicates, mean + standard deviation).

Subsequently, growth of the pTAT_cellulase P_*EM7*_-cbh strain was evaluated in minimal media with glucose, cellotriose, or cellotetraose as sole carbon source (Figure 4), without addition of 3-methylbenzoic acid. The maximum specific growth rate of pTAT_cellulase P_*EM7*_-cbh grown on glucose (0.31 h^-1^ ± 0.02) was comparable to the pSEVA231 control strain (0.33 h^-1^ ± 0.02), though a growth penalty for the cellulase secretion strain was observable in the linear growth phase (Figure 3A). pTAT_cellulase P_*EM7*_-cbh grew with cellotriose or cellotetraose as the sole carbon source but with significantly slower growth rates, longer lag phases, and lower total biomass accumulation (Figure 4, Table 2).

**Table 2.** Growth parameters on glucose, cellotriose, and cellotetraose.

| Carbon source | Strain | $\mu$ ($h^{-1}$) | Lag phase (h) | Maximum biomass (OD<sub>600 nm</sub>) |
| --- | --- | --- | --- | --- |
| Glucose | pTAT_cellulase $P_{EM7\_cbh}$ | $0.31 \pm 0.02$ | $6.3 \pm 0.5$ | $1.07 \pm 0.01$ |
| | pSEVA231 $P_{EM7\_cbh}$ | $0.34 \pm 0.02$ | $5.7 \pm 0.5$ | $1.11 \pm 0.05$ |
| Cellotriose | pTAT_cellulase $P_{EM7\_cbh}$ | $0.01 \pm 0.01$ | $113 \pm 16$ | $0.67 \pm 0.12$ |
| | pSEVA231 $P_{EM7\_cbh}$ | - | - | $0.26 \pm 0.01$ |
| Cellotetraose | pTAT_cellulase $P_{EM7\_cbh}$ | $0.01 \pm 0.01$ | $145 \pm 32$ | $0.52 \pm 0.13$ |
| | pSEVA231 $P_{EM7\_cbh}$ | - | - | $0.26 \pm 0.01$ |

$n = 3$ biological replicates ± standard deviation

**Figure 4.**
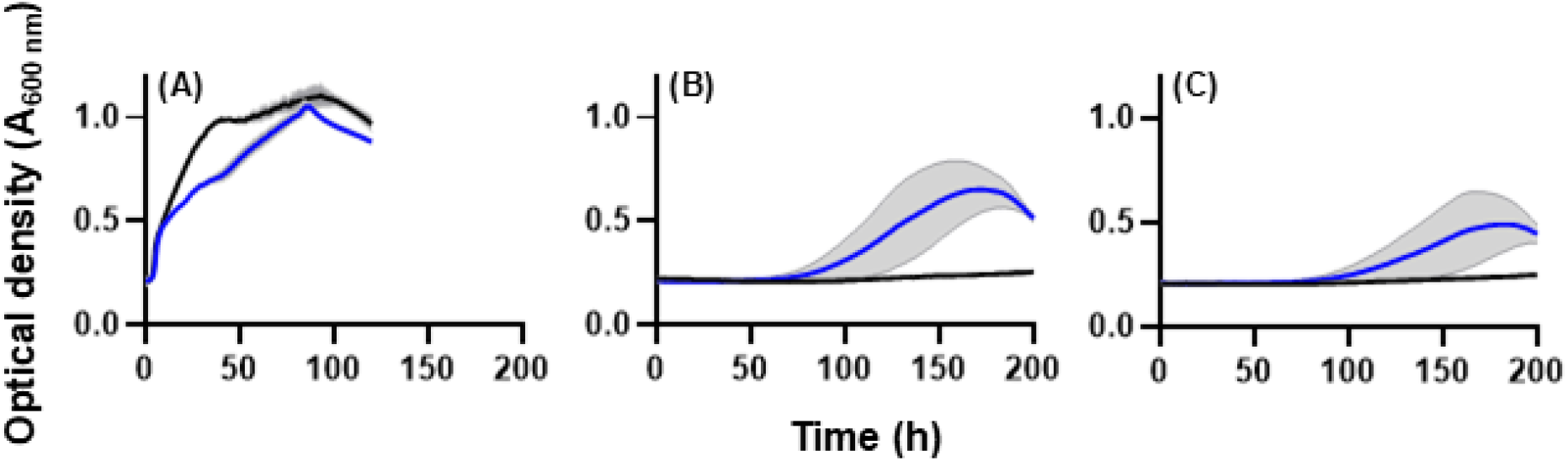
Growth with glucose, cellotriose or cellotetraose as sole carbon source. *P. putida* strains pTAT_cellulase P_*EM7*__cbh (blue line) or *P. putida* pSEVA231 P_*EM7*__cbh (black line) were grown in defined minimal medium liquid culture with (A) glucose, (B) cellotriose, or (C) cellotetraose as the sole carbon source (200 μL volumes in a 96-well microtitre plate). Growth (measured as optical density at 600 nm) was monitored for 200 h (n = 3 biological replicates, plots display mean optical density ± standard deviation shown with grey shading).

No growth was observed when cellulase secretion strains were incubated in liquid medium containing crystalline cellulose as the sole carbon source, whether in the form of cellulose filter paper or microcrystalline Avicel. Likewise, significant growth was not observed when regenerated amorphous cellulose or phosphoric acid swollen cellulose was supplied as the sole carbon source in solid agar medium.

## Discussion

These data demonstrate secretion of cellulase enzymes by an engineered *P. putida* strain and show that extracellular cellulase activity is sufficient to support growth using oligomeric cellulose substrates (cellotriose or cellotetraose) as the sole source of carbon and energy, though at significant fitness cost relative to growth on glucose. This is a new development in the engineering of *P. putida* for versatile carbon source utilisation, expanding on previous work where *P. putida* strains were modified for intracellular metabolism of xylose, cellobiose, sucrose, and arabinose (Meijnen et al. 2008; Löwe et al. 2017; Dvořák and de Lorenzo 2018; Bator et al. 2020).

Meaningful growth on cellulose oligomers was only observed when cellulases were secreted using the signal peptides encoded on the pTAT_cellulase plasmid and co-expressed with an intracellular cellobiohydrolase. Growth on cellotriose was slow, with a maximum specific growth rate less than 10% of that observed for growth on glucose. The lag phase was prolonged by approximately four days, and increased by a further 28% when grown on cellotetraose. The low growth rates and reduced biomass titres indicate a sub-optimal metabolism where there is insufficient extracellular cellulase activity to support high growth rates.

Extracellular cellulase activity could not be improved simply by increasing protein translation; growth of the pTAT_cellulase P_*EM7*_-cbh strain on cellotriose was significantly worse when gene expression was induced by addition of 3-methylbenzoic acid. The xylS/Pm promoter system is known to support relatively high levels of background expression in the uninduced state (Volke et al. 2020), and uninduced expression from the pTAT_cellulase plasmid was sufficient to support growth on cellulose oligomers.

Toxicity caused by overexpression of secreted proteins and saturation or blockage of their secretion pathways has been recognised for almost fifty years (Bassford and Beckwith 1979). Saturation of Sec or Tat secretion pathways is a limiting factor in heterologous protein secretion (Ignatova et al. 2003; DeLisa et al. 2004), and successful secretion of heterologous proteins in gram negative bacteria requires careful calibration of expression levels (Schlegel et al. 2013) to balance synthesis with capacity for export. The cellulase secretion phenotype could potentially be optimised via adaptive laboratory evolution, where protein synthesis and capacity for secretion could be evolved in tandem by selecting for increased growth rates of strains grown in the presence of cellulose oligomers of increasing molecular weight. Previous studies observed that saturation of the Tat pathway secretion can be alleviated by upregulation of Tat complex expression and membrane stress response proteins (such as the phage shock protein pspA), and that this in turn increases the yield of secreted proteins (DeLisa et al. 2004; Matos et al. 2012). Sec (Natarajan et al. 2017) and Tat (Taw et al. 2021) translocase activity were recently evolved in *E. coli* for enhanced rates of secretion by selecting for increased secretion of a beta-lactamase enzyme into the periplasmic space (selected by exposure to increasing concentrations of beta-lactam antibiotics). Enhancement of extracellular secretion (across the outer membrane) was achieved by co-expression of a beta-lactamase inhibitor protein (BLIP) fused to a YebF extracellular secretion peptide (Natarajan et al. 2017). Accumulation of the BLIP in the periplasm conferred antibiotic sensitivity, which was relieved by enhanced secretion of the YebF-BLIP fusion to the extracellular environment. Growth using polymeric cellulosic substrates is promising as a selective pressure for evolving two-step translocation of active cellulases, and potentially other heterologous proteins, to the extracellular environment of *P. putida*.

Cellulase activity in the extracellular fraction was consistently greatest when using the selected Tat pathway secretion signals, though it is not clear whether this is due to differences in translocation efficiency with the selected signal peptides, differences in compatibility between the cellulase cargo proteins and the Sec or Tat secretion pathways, or differences in transcription and translation. The Sec pathway translocates proteins across the inner membrane in an unfolded state, whereas the Tat pathway transports folded proteins via a pore formed by the TatABC complex. Physical properties of the cargo protein, including folding dynamics, are important determinants of translocase compatibility (Kleiner-Grote et al. 2018). Mechanisms underlying the second translocation step from the periplasm across the outer membrane remain poorly understood even in *E. coli* and there are few data sets available from which extracellular proteins and their secretion signal peptides can be identified (Qian et al. 2008; Salvachúa et al. 2020).

*P. putida* is particularly suited to manufacturing commodity chemicals (Weimer et al. 2020) like acids and alcohols, which attract a low selling price. Therefore accessing suitably low-cost feedstocks is essential to developing cost-effective manufacturing processes that can compete with equivalent fossil-derived products (Werner et al. 2023). Expanding the range of feedstocks that *P. putida* can utilize offers the chance to reduce fermentation costs by making better use of fermentable sugars, and establishment of a strain capable of secreting hydrolytic enzymes expands the design possibilities for reducing bioprocessing costs in microbial fermentations (Raftery and Karim 2017; Dempfle et al. 2021). In addition to development of conventional single-strain consolidated bioprocessing, it may be possible to build on the recent expansion of sugars accessible by engineered *P. putida* (Meijnen et al. 2008; Löwe et al. 2017; Dvořák and de Lorenzo 2018; Bator et al. 2020) to develop subpopulations with distinct carbohydrate preferences and bioprocessing roles.

## Materials and Methods

### Chemicals, microbial strains, and culture media

Congo Red was obtained from Merck (catalogue no. C6277), cellotriose was obtained from Neogen (http://megazyme.com, product code O-CTR-50MG), Avicel PH-101 was obtained from Merck (catalogue no. 11365), 3-methylbenzoic acid was obtained from Merck (catalogue no. 8.21902).

M9 medium in this instance refers to final concentrations of disodium phosphate (47 mM), monopotassium phosphate (22 mM), sodium chloride (0.5 g/L), ammonium chloride (1 g/L), and magnesium sulfate (2 mM). In all instances, Studier’s trace metals(Studier 2005) were included at a 1x concentration. The trace metal mixture supplies the calcium chloride ordinarily included in an M9 medium, as well as other elements in which M9 is deficient.

### Genetic designs

Genetic constructs are described in Table 1 and Online Resource 1 (Supporting data S1). Briefly, the native secretion signal peptides for the CelA endocellulase (Genbank ID: WP_003512420.1) and CelK exocellulase (Genbank ID WP_011837826.1) from *Acetivibrio thermocellus*(Tozakidis et al. 2016) were predicted using SignalP 6.0(Teufel et al. 2022). The native signal peptide was replaced by in-frame fusion with a *P. putida* Sec or Tat signal peptide identified in the literature as being associated with extracellular *P. putida* proteins (Table 1). CelA and CelK constructs were arranged in an operon, and *P. putida* codon optimization and translation initiation rate (TIR) for each gene were designed using the Operon Calculator from DeNovo DNA (Farasat et al. 2014) (http://www.denovodna.com). Designs were synthesised by Twist Bioscience (http://twistbioscience.com) and cloned into a pSEVA231 (Martínez-García et al. 2023) expression vector under the control of the xylS/Pm (Gawin et al. 2017) inducible promoter.

To create the *P. putida* KT2440_P_*EM7*_-cbh strain (constitutive intracellular β-glucosidase expression), we used a previously-published (Dvořák and de Lorenzo 2018) genetic design where cellobiose hydrolysis was encoded by the *bglC* gene of *Thermobifida fusca* (cbh; Genbank accession WP_011291384.1) under the control of the constitutive P_*EM7*_ promoter. The P_*EM7*_-cbh construct was synthesised as a linear fragment (Integrated DNA Technologies, http://www.idtdna.com) and cloned into the pBAMD1-6 mini-Tn5 transposon delivery plasmid (Martínez-García et al. 2014) using the general principles of Gibson isothermal assembly (Gibson et al. 2009). Transformed *P. putida* were subject to an extended post-transformation recovery in terrific broth (7 h, 30 °C, 180 rpm agitation)

The total recovered cell population was washed and resuspended in 100 mL M9 medium supplemented with 5 g cellobiose/L, 30 mg gentamycin/L, and Studier’s trace metals (Studier 2005), and incubated for 4 days (30 °C, 180 rpm agitation) before plating on M9 agar supplemented with 5 g cellobiose/L, 30 mg gentamycin/L, and Studier’s trace metals. Gene integration was confirmed by colony PCR and Sanger sequencing. Growth performance with glucose or cellobiose as sole carbon source was compared for 5 individual clones. Individual transformant clones were pre-cultured in M9 minimal medium containing glucose (5 g/L) or cellobiose (5 g/L), 30 °C, 180 rpm shaking in a standard orbital shaker. Untransformed *P. putida* KT2440 was included as a control. Starter cultures inoculated at a 1 in 100 dilution into 200 µL M9 minimal medium containing either glucose (5 g/L, black lines) or cellobiose (5 g/L, blue lines), with n = 4 technical replicate cultures per clone, and cultivated at 30 °C with 900 rpm shaking and monitoring absorbance at 600 nm every ten minutes. All cultures included gentamycin (30 mg/L) except for those of untransformed *P. putida* KT2440. Colony 8, which showed the best growth performance with cellobiose as sole carbon source (Supporting Figure S6), was selected as a single *P. putida* KT2440_P_*EM7*_-cbh strain for subsequent transformation with cellulase secretion plasmids.

Cellulase secretion plasmids (pSEC_cellulase and pTAT_cellulase) were used to transform *P. putida* KT2440_P_*EM7*_-cbh, *P. putida* KT2440, or *P. putida* S12. Plasmids were maintained by selection with kanamycin (50 mg/L). Negative control strains were prepared by transforming with the pSEVA231 vector. All experiments used three biological replicates for each strain, where a biological replicate is defined as an independent colony from the plasmid transformation step.

### Carboxymethylcellulose hydrolysis on solid media

Strains were pre-cultured overnight (30 °C with agitation) in LB medium supplemented with 50 μg kanamycin/mL (plus 30 μg gentamycin/mL for strains in the *P. putida* KT2440_cbh background). Overnight cultures were washed in unsupplemented M9 medium and diluted to OD_600 nm_ = 0.3. 5 μL volumes of each culture were spotted onto M9 agar supplemented with antibiotics, Studier’s trace metals, yeast extract (0.05% w/v), carboxymethylcellulose (CMC: 0.5% w/v), and 3-methylbenzoate (1 mM). Agar plates were incubated at 30 °C for 64 h. CMC hydrolysis was observed following CMC staining with Congo Red (Teather and Wood 1982) (15 min incubation with 5 mL of Congo Red, 1 mg/mL, followed by 15 min washing with 5 mL sodium chloride, 1 M).

### Cellulase enzyme activity assays

#### Assay pre-culture

Strains were pre-cultured overnight (30 °C, 180 rpm) in M9 medium supplemented with 0.2% (w/v) casamino acids and 50 μg kanamycin/mL (and 30 μg gentamycin/mL for strains in the *P. putida* KT2440_cbh background). Overnight cultures were diluted to OD600 nm = 0.05 in 1 mL M9 media supplemented with 0.2% (w/v) casamino acids. After 3 h, cellulase expression was induced by addition of 1 mM 3-methylbenzoic acid then incubated overnight (16 h, 30 °C, 180 rpm). Cells were sedimented via centrifugation (5 min at 3900 rpm) and culture supernatants were removed and assayed to determine the presence of cellulases (detailed below).

#### 4-hydroxybenzoic acid hydrazide (PAHBAH) assay

The 4-hydroxybenzoic acid hydrazide (PAHBAH) assay for detection of reducing sugar ends(Lever 1972) was adapted from previously published methods (Garvey et al. 2014).

Carboxymethylcellulose was added to culture supernatants at a final concentration of 1% (w/v) and incubated at 50 °C with agitation for 4 h. Samples were taken hourly and immediately combined with PAHBAH working reagent and incubated at 95 °C for 10 minutes and then placed on ice for 5 minutes. Sample volumes were 30 μL combined with 200 μL working reagent that contained 36 mM PAHBAH. Post-incubation, absorbance at 405 nm was measured in a ThermoFisher Varioskan multimode plate reader.

#### 4-methylumbelliferyl-β-D-cellobioside (4-MUC) assay

4-methylumbelliferyl-β-D-cellobioside (4-MUC) was added to culture supernatant samples to a final concentration of 1.25 mM and incubated for 3 h at 50 °C. Reactions were quenched by addition of 100 μL 0.1 M sodium carbonate, and diluted 1 in 10 in water before fluorescence was measured (λex = 360 nm, λex = 460 nm) in a ThermoFisher Varioskan multimode plate reader.

#### 4-Nitrophenyl β-D-cellobioside (p-NPC) assay

4-Nitrophenyl β-D-cellobioside (*p*-NPC) was added to culture supernatant samples to a final concentration of 0.5 mM and incubated for 2 h at 45 °C. After 2 h, absorbance at 405 nm was measured in a ThermoFisher Varioskan multimode plate reader.

### Proteomic identification of cellulases

Targeted mass spectrometry-based proteomic analysis was used to qualitatively identify heterologous expression of endocellulase (CelA) and exocellulases (CelK) in *P. putida* culture supernatants and cell pellets. Cultures grown in LB medium were harvested in late exponential phase, and supernatant and cell pellet fractions were separated by centrifugation at 4,500 *g*, 15 min, 4 °C. Supernatants were concentrated 20-fold using a 10 kDa molecular weight cutoff filter (Vivaspin 20 Cat. No. 28-9322-60). Cell pellet or concentrated supernatant equivalent to 100 μg total protein were mixed with 100 μL lysis buffer (4% SDS, 100 mM Tris-HCl (pH 8), 100 mM DTT) and heated at 95 °C for 5 min. Cell pellet samples were lysed by bead beating with acid washed glass beads (Sigma Cat. No. G1152) at 30 s^-1^ for using a Qiagen Tissue Lyser II, followed by brief ultrasonication. The insoluble fraction was sedimented by centrifugation (20,000 *g*, 20 min). Cell lysates were sonicated with ultrasound for a few pulses and pelleted by centrifugation. Cell lysates were then digested with trypsin according to the Filter Aided Sample Preparation protocol (FASP) (Wiśniewski et al. 2009) and purified with C-18 tips (Pierce C18 Pippete Tips Cat. No. P187784). Purified samples were vacuum dried and resuspended in iRT buffer (0.1% Formic Acid, iRT Kit Biognosys Cat. No. Ki-30002-1) (Escher et al. 2012). Trypsin digested peptides of samples were MS analysed in Data Dependent Acquisition mode (DDA) using the Sciex 5600 QTOF. Protein pilot 5.0.2, using the Paragon method and amino acid sequence of engineered cellulases, was applied to search and identify peptides in the samples at a quality threshold of 1.3 (95%). Skyline-daily 23.1.1.425 was used to confirm the quality control of injections and trypsin digestion by identifying the retention time of iRT peptides and BSA trypsin digested sample.

### Growth of *P. putida* on cellulose oligomers

*P. putida* strains were precultured in M9 mineral medium supplemented with glucose (0.5 %, w/v), Studier’s trace metals (Studier 2005) and 50 μg kanamycin/mL. Seed cultures were washed twice with M9 salts containing no carbon source and inoculated into fresh M9 medium supplemented with trace metals and kanamycin (50 µg/mL), and either 0.5 % glucose or 0.5 % cellotriose (w/v). Cultures were grown in 200 μL volumes in a standard 96-well microtitre plate sealed with a breathe-easy membrane (Merck catalogue no. Z380059). In preliminary screening of pSEC_cellulase and pTAT_cellulase P_*EM7*__cbh strains grown on glucose and cellotriose, cultures were incubated in a ThermoFisher Varioskan multimode plate reader (30 °C, 600 rpm agitation) and absorbance at 600 nm was measured every 30 minutes for 120 hours. In a subsequent experiment, pTAT_cellulase P_*EM7*__cbh and pSEVA_231 P_*EM7*__cbh strains were inoculated to an initial OD_600 nm_ of 0.2 in supplemented M9 medium with either 0.5 % glucose, 0.5 % cellotriose, or 0.5 % cellotetraose (w/v) as sole carbon source, and agitation was increased to 900 rpm and absorbance was monitored for 200 hours.

### Assessment of growth on high molecular weight cellulose in solid medium

Regenerated amorphous cellulose (RAC) was prepared as described previously (Rinaldi 2011) with the following modifications: 10 g of cellulose was combined with 80 g of DMSO and stirred at 100 °C for 20 minutes at 300 rpm. Upon addition of 20 g of 1-ethyl-3-methylimidazolium acetate, the cellulose rapidly dissolved. The cellulose solution was then regenerated by adding it to 300 mL of cold water. The resulting regenerated cellulose was thoroughly washed with water, freeze-dried, and ground to a fine powder. The crystallinity index of RAC, assessed through quantitative X-ray powder diffraction (XRD), indicated a composition with 92.3% amorphous content and 7.7% crystalline content (Online Resource 1, Supporting Methods S1 and Supporting Figure S8). Phosphoric acid swollen cellulose (PASC) was prepared as described previously (Den Haan et al. 2007). RAC or PASC were added to M9 agar (M9 salts plus Studier’s trace metals and 50 μg kanamycin/mL) at a concentration of 0.5 % (w/v). Strains were grown overnight in LB medium supplemented with kanamycin (50 μg/mL) and then streaked onto M9+PASC or M9+RAC agar. Replicate plates were incubated at 30 °C and 40 °C for up to 21 days. Strains were also streaked onto M9 agar with glucose (0.5% w/v) or M9 agar with no carbon source as positive and negative controls, respectively.

## Supporting information

Online Resource 1

## Acknowledgements

The pSEVA231 plasmid was a generous gift from the Standard European Vector Architecture collection (https://seva-plasmids.com). Proteomic and X-ray powder diffraction data reported in this paper were obtained at the Central Analytical Research Facility operated by Research Infrastructure (QUT).

## Author Contributions

M. S., K. M., and W. H. conducted the microbial growth experiments and enzyme assays. C. L-F. performed the proteomic analyses. M. G. prepared and analysed the regenerated amorphous cellulose. M. S., R. S., A. B. and J. B. developed the concept and designed experiments. All authors contributed to writing and revision of the manuscript.

## Funding Declaration

This project was supported by a Masters Scholarship awarded to M. R. Smith by Sugar Research Australia (Project number 2020/101) and a research fellowship awarded to J. Behrendorff by the CSIRO Synthetic Biology Future Science Platform. This work was also supported by the Australian Research Council Centre of Excellence in Synthetic Biology (project CE200100029).

## Conflict of interest statement

The authors declare no conflicts of interest.

## Data availability statement

The authors declare that the data supporting the findings of this study are available within the paper and its Supplementary Information files. Should any raw data files be needed in another format they are available from the corresponding author upon reasonable request.

## Ethics declaration

not applicable

## Notes

### Competing Interest Statement

The authors have declared no competing interest.

