## Supplementary material for "Cellulase secretion by engineered *Pseudomonas putida* enables growth on cellulose oligomers": Online Resource 1

**Supporting Table S1 Extracellular protein secretion signals identified in published literature.**

| Source protein (Genbank accession) | Secretion pathway (secretion signal residues adapted for this study) | Description | Use in this study |
|---|---|---|---|
| uxpB (WP_284383961.1) | TAT (amino acids 1-59, MSRDTGDNLDRNQSGNLPMANVMDAYLSRRSVMRGSLGAAIAMIAGTGLTGCFDGGGSD) | Extracellular phosphatase natively produced by *P. putida* under low phosphate conditions[1] | pTAT_cellulase (secretion of CelK)<br>pUxpB_celK (secretion of CelK)<br>pUxpB_celA_1K (secretion of CelA, TIR 1000)<br>pUxpB_celA_10k (secretion of CelA, TIR 10,000) |
| PP_2478 (WP_010953415.1) | TAT (amino acids 1-45), MKKPNEVTVDMSRRRLLQGSGIALSGLVLSTWLPPLVAKSAAAEA | Isoquinoline 1-oxidoreductase, beta subunit. Identified exclusively in *P. putida* the extracellular fraction in a proteomic study of outer membrane vesicles[2]. | pTAT_cellulase (secretion of CelA)<br>p2478_celA_1K (secretion of CelA, TIR 1000)<br>p2478_celA_10k (secretion of CelA, TIR 10,000) |
| OprF | Sec (amino acids 1-24, MKLKNTLGLAIGSLVAATSIGAMA) | Porin F, major outer membrane protein of *P. putida* abundant in outer membrane vesicles[3] | pSEC_cellulase (secretion of CelK) |
| PP_5130 (WP_010955666.1) | Sec (amino acids 1-21, MKVAVKAAAIGLSLLFSIETF) | Phosphorylcholine phosphatase identified exclusively in *P. putida* the extracellular fraction in a proteomic study of outer membrane vesicles[2]. | pSEC_cellulase (secretion of CelA) |

**Supporting Table S2 Proteomic identification of cellulases in culture supernatant**

| Strain | Fraction | CelK peptides detected (>95% confidence) | CelA peptides detected (>95% confidence) |
|---|---|---|---|
| pTAT_cellulase | Supernatant | YLRPVSTAATLNFAATLAQSAR<br>YLDGMQDGMSYLLGR | GIVDGYTIQGSK<br>MKKPNEVTVDMSR |
| | Cell pellet | YLRPVSTAATLNFAATLAQSAR<br>FDALAFFYHKR | Not detected |
| pSEC_cellulase | Supernatant | YLRPVSTAATLNFAATLAQSAR<br>YLDGMQDGMSYLLGR | GIVDGYTIQGSK<br>MKKPNEVTVDMSR |
| | Cell pellet | YLRPVSTAATLNFAATLAQSAR<br>FDALAFFYHKR | Not detected |

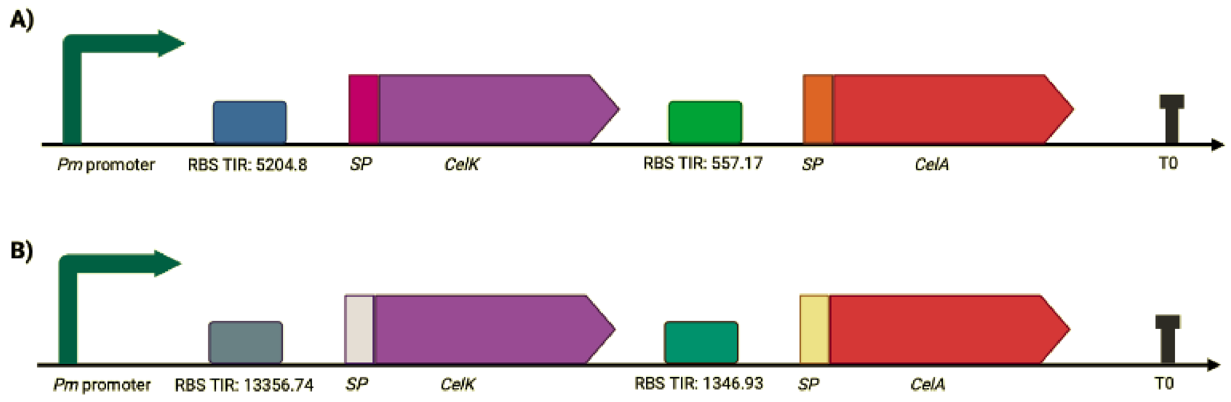


**Supporting Figure S1 Schematic of genetic constructs for cellulase secretion.** A) pSEC_cellulase operon design: Pm promoter (xylS/Pm inducible system), ribosome binding site (RBS) with designed translation initiation rate (TIR) of 5204.8 for the CelK gene bearing a Sec pathway secretion signal fused in-frame to its 5' end, RBS with TIR 557.17 for the CelA gene bearing a Sec pathway secretion signal fused in-frame to its 5' end, T0 transcriptional terminator. B) pTAT_cellulase operon design: Pm promoter, RBS with TIR of 13356.74 for CelK with a Tat pathway secretion signal, RBS with TIR 1346.93 for CelA gene with a Tat pathway secretion signal, T0 transcriptional terminator. Secretion signal peptides are detailed in Supporting Tabel 1, and full coding sequences are detailed in Supporting Data 1. Synthetic expression operons were designed at http://www.denovodna.com.

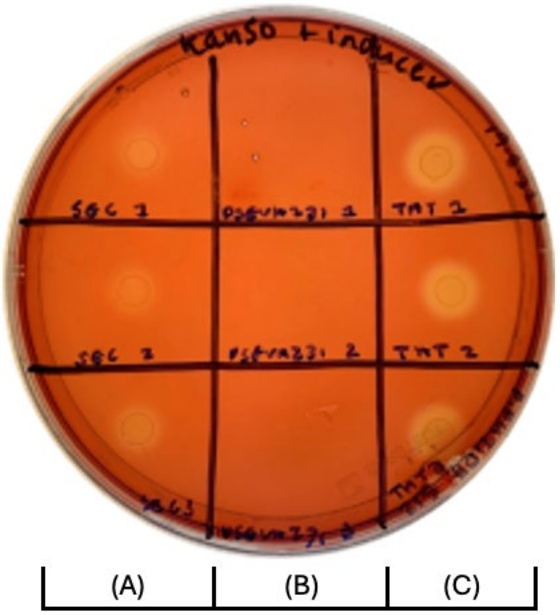


**Supporting Figure S2 Carboxymethylcellulose hydrolysis on solid agar.** Liquid cultures (5 µL) of *P. putida* S12 transformed with pSEC_cellulase (A), pSEVA231 (B), and pTAT_cellulase (C) plasmids were spotted on M9 agar containing carboxymethylcellulose (0.5% w/v). Carboxymethylcellulose hydrolysis was visualised by staining with Congo Red. Three biological replicates for each strain are arranged in each marked column.

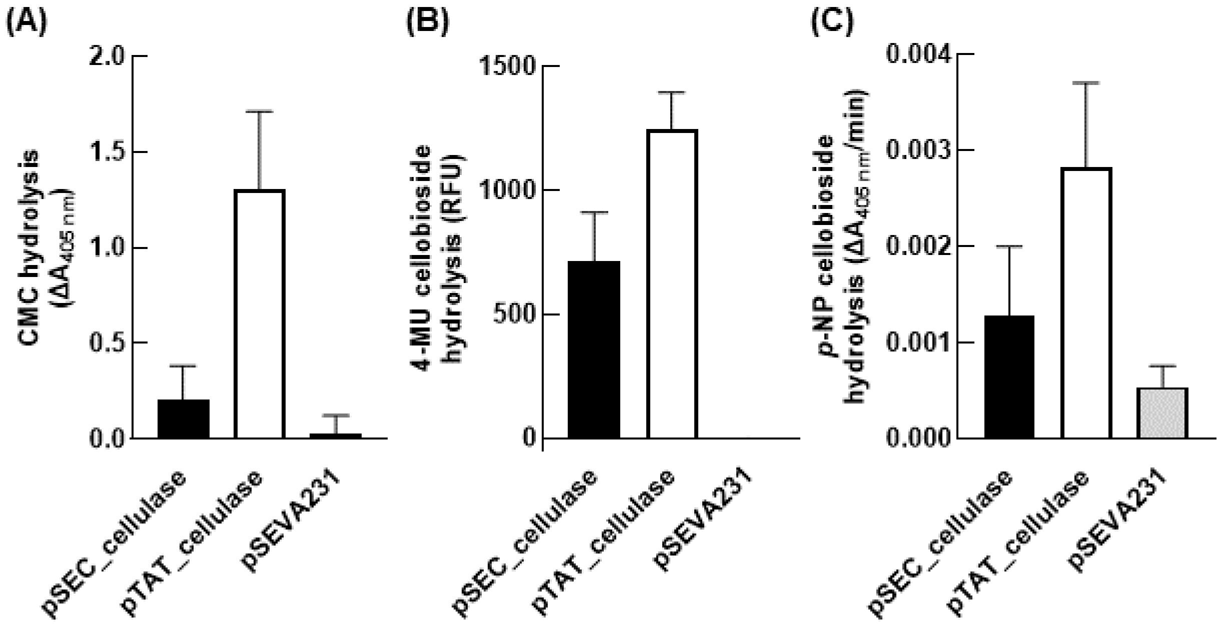


**Supporting Figure S3 Cellulase activity in *P. putida* S12 culture supernatants.** *P. putida* S12 expression cultures were sedimented via centrifugation and the culture supernatants were incubated with a variety of probe substrates for detection of cellulase activity. (A) Carboxymethylcellulose (CMC) hydrolysis was assayed via reaction of reducing sugar ends with *p*-hydroxybenzoic acid hydrazide. (B) 4-methylumbelliferyl β-D-cellobioside (4-MU cellobioside) hydrolysis was assayed as fluorescence of liberated 4-methylumbelliferone. (C) *p*-nitrophenyl β-D-cellobioside (*p*-NP cellobioside) hydrolysis was assayed as change in absorbance from liberated *p*-nitrophenol, comparing rate of product formation in the initial linear phase of the reaction. All plots report the mean of n=3 biological replicate cultures ± standard deviation.

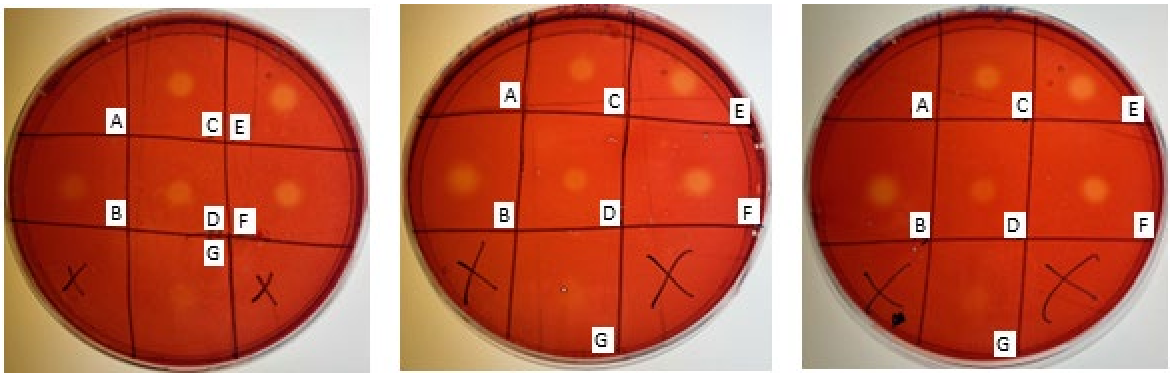


**Supporting Figure S4 Carboxymethylcellulose hydrolysis by individual cellulases.** Liquid cultures (5 µL) of *P. putida* KT2440 transformed with pSEVA231 (A), pTAT_cellulase (B), p2478-celA_1k (C), pUxpB-celA_1K (D), pUxpB-celA_10k (E), p2478-celA_10K (F), and pUxpB-celK (G) plasmids were spotted on M9 agar containing carboxymethylcellulose (0.5% w/v). Carboxymethylcellulose hydrolysis was visualised by staining with Congo Red. Three biological replicates for each strain are displayed on separate agar plates.

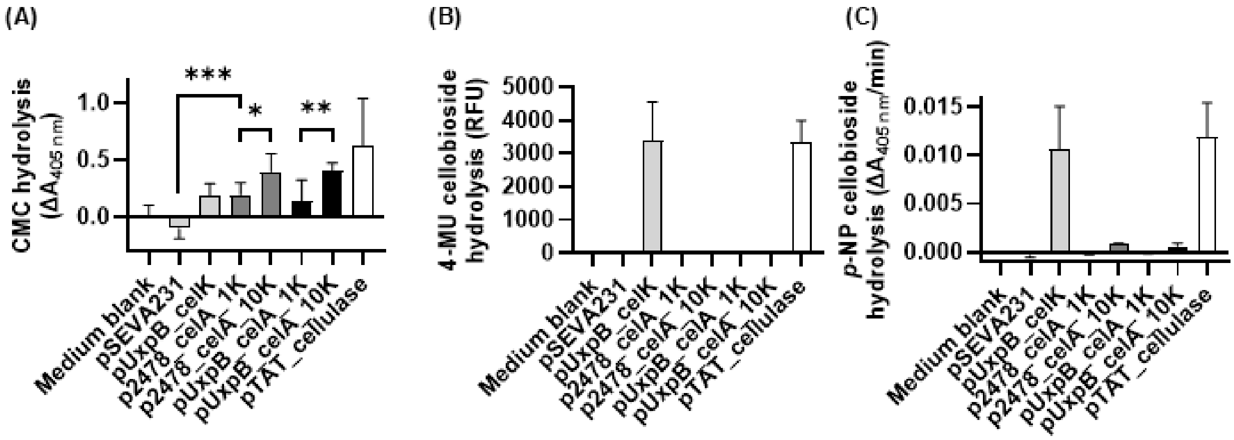


**Supporting Figure S5 Cellulase activity in culture supernatants.** *P. putida* KT2440 expression cultures were sedimented via centrifugation and the culture supernatants were incubated with a variety of probe substrates for detection of cellulase activity. (A) Carboxymethylcellulose (CMC) hydrolysis was assayed via reaction of reducing sugar ends with *p*-hydroxybenzoic acid hydrazide. (B) 4-methylumbelliferyl β-D-cellobioside (4-MU cellobioside) hydrolysis was assayed as fluorescence of liberated 4-methylumbelliferone. (C) *p*-nitrophenyl β-D-cellobioside (*p*-NP cellobioside) hydrolysis was assayed as change in absorbance from liberated *p*-nitrophenol, comparing rate of product formation in the initial linear phase of the reaction. All plots report the mean of n = 3 biological replicate cultures + standard deviation. unpaired Student's *t*-test (* = $p < 0.05$, ** = $p < 0.01$, *** = $p < 0.001$).

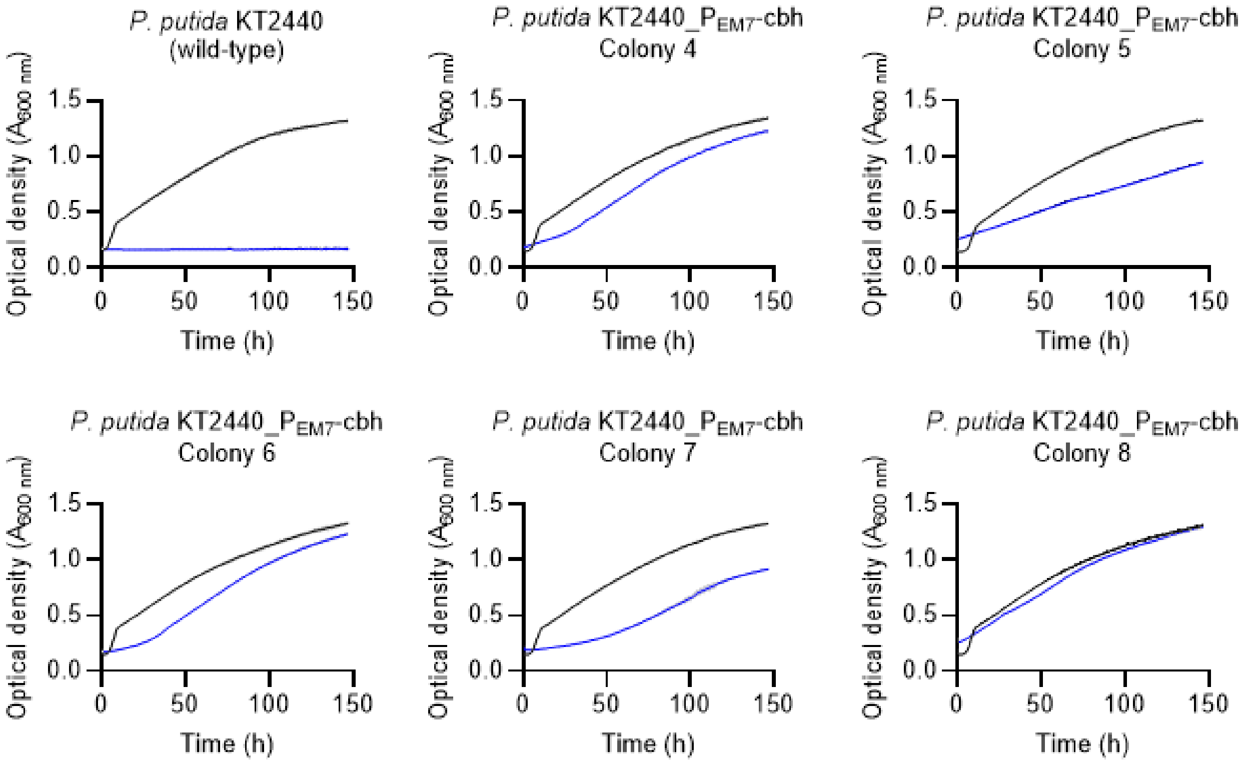


**Supporting Figure S6 Selection of a cellobiose-metabolising strain, *P. putida* KT2440_P$_{EM7}$-cbh.** *P. putida* KT2440 was transformed with a β-glucosidase under the control of the constitutive P$_{EM7}$ promoter. Five transformant colonies were identified via colony PCR and DNA sequencing of the transgene, and evaluated for growth on glucose (black lines) and cellobiose (blue lines). Individual transformant clones were pre-cultured in M9 minimal medium containing glucose (5 g/L) or cellobiose (5 g/L). Starter cultures inoculated at a 1 in 100 diultion into M9 minimal medium containing either glucose (5 g/L, black lines) or cellobiose (5 g/L, blue lines), with n = 4 technical replicate cultures per clone. All cultures included gentamycin (30 mg/L) except for those of untransformed *P. putida* KT2440. Curves display mean of replicate cultures ± standard deviation (in grey shading either side of curve).

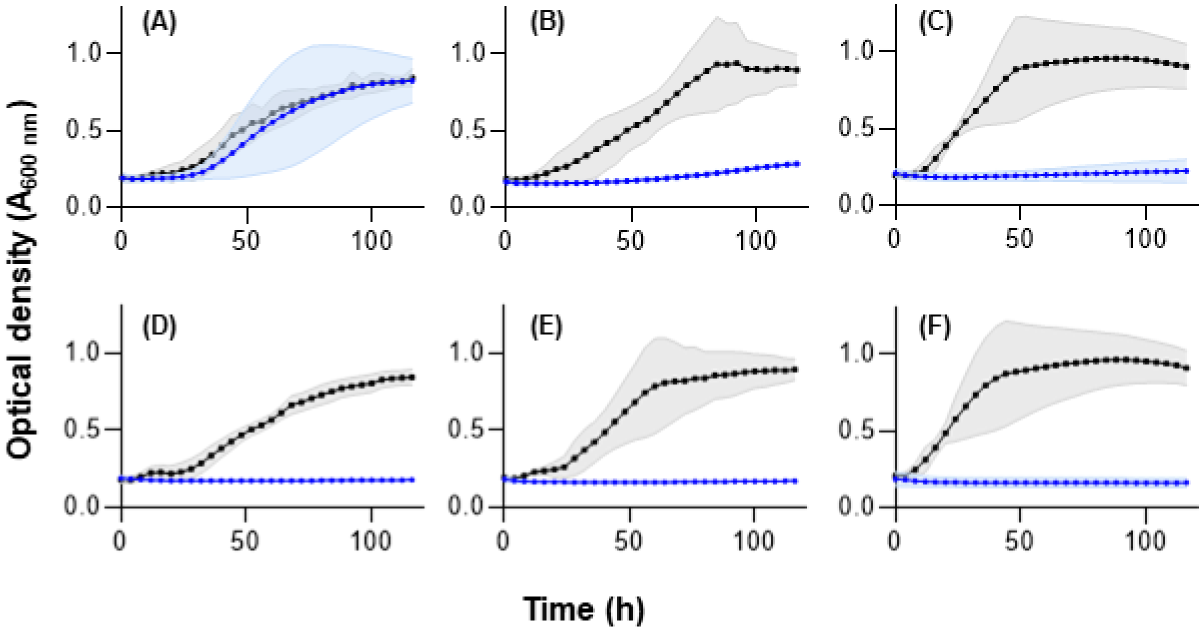


**Supporting Figure S7 Growth of cellulase secretion strains with cellotriose as sole carbon source.** *P. putida* strains were grown in defined minimal medium liquid cultures with cellotriose (blue) or glucose (black) as the sole carbon source (200 µL volumes in a 96-well microtitre plate, 30 °C, 600 rpm continuous shaking). (A) pTAT_cellulase $P_{EM7}$_cbh, (B) pSEC_cellulase $P_{EM7}$_cbh, (C) pSEVA231 $P_{EM7}$_cbh, (D) pTAT_cellulase, (E) pSEC_cellulase, (F) pSEVA231. Growth was monitored for 120 h (n = 3 biological replicates, plots display mean optical density ± standard deviation shown with shading).

**Supporting Method S1**

X-ray powder Diffraction (XRD) was used to estimate the crystallinity index (CrI) of the cellulose sample. XRD patterns were acquired using a Bruker D8 Advance powder diffractometer operating in Bragg-Brentano geometry with a cobalt source (35 kV, 40 mA). Patterns were collected for 60 minutes from 2 to 89 °2θ at a step size and of 0.015°. Samples were spun during data collection at spun rate of 15rpm. Incident optics included 2.5° Soller slits, and a variable divergence slit with an illuminated length of 10 mm. Receiving optics before the LYNXEYE XE-T detector (high resolution mode) included 2.5° Soller slits, and an open (18 mm) receiving slit. The D8 Advance had a goniometer radius of 280 mm, and the detector had an opening of 2.945°. An automatic beam knife provided superior background suppression and reduced air scatter at low angles.

The crystallinity index of cellulose sample was calculated based on the Segal method(Segal et al. 1959) i.e., from the height ratio of intensity of crystalline peak (I200- I¬AM) and total intensity (I200) excluding background signal. The phase identification was performed using PDF4+ database (ICDD, 2021) in EVA (V5, Bruker). The Rietveld method as implemented in TOPAS (V7, Bruker) was used for refinement and quantitative phase analysis in which the crystalline component was calculated from the structure of cellulose Iβ. The degree of crystallinity was determined to be 7.7% by calculating the ratio of area of crystalline cellulose and the total area of crystalline plus amorphous cellulose.

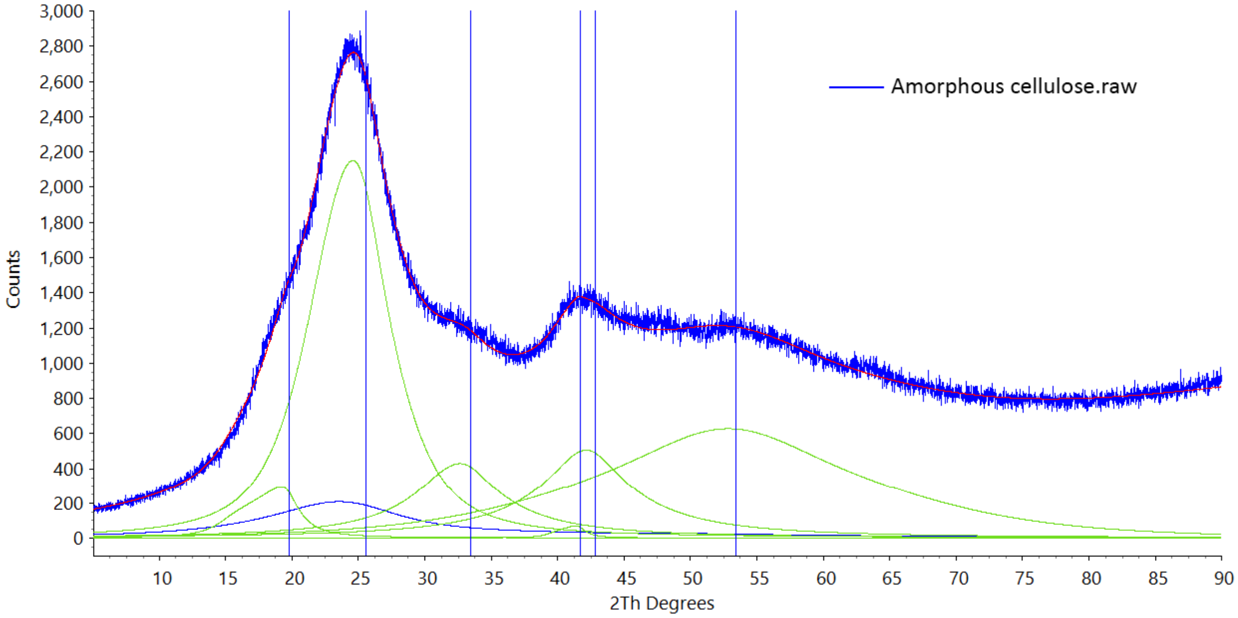


**Supporting Figure S8** X-ray powder diffraction trace of regenerated amorphous cellulose**.**

**Supporting data S1 Cellulase sequence information**

Protein sequences of cellulases from the pSEC_cellulase and pTAT_cellulase plasmids used in this study, including in-frame fusion to Sec or Tat signal peptides (underlined).

>SEC_celA (from pSEC_cellulase)

MKVAVKAAAIGLSLLFSIETFAGVPFNTKYPYGPTSIADNQSEVTAMLKAEWEDWKSKRITSNGAGGYKRVQRDASTNY
DTVSEGMGYGLLLAVCFNEQALFDDLYRYVKSHFNGNGLMHWHIDANNNVTSHDGGDGAATDADEDIALALIFADKL
WGSSGAINYGQEARTLINNLYNHCVEHGSYVLKPGDRWGGSSVTNPSYFAPAWYKVYAQYTGDTRWNQVADKCYQI
VEEVKKYNNGTGLVPDWCTASGTPASGQSYDYKYDATRYGWRTAVDYSWFGDQRAKANCDMLTKFFARDGAKGIV
DGYTIQGSKISNNHNASFIGPVAAASMTGYDLNFAKELYRETVAVKDSEYYGYYGNSLRLLTLLYITGNFPNPLSDLSGQP
TPPSNPTPSLPPQVVYGDVNGDGNVNSTDLTMLKRYLLKSVTNINREAADVNRDGAINSSDMTILKRYLIKSIPHLPY

>TAT_celA (form pTAT_cellulase)

MKKPNEVTVDMSRRRLLQGSGIALSGLVLSTWLPPLVAKSAAAEAAGVPFNTKYPYGPTSIADNQSEVTAMLKAEWED
WKSKRITSNGAGGYKRVQRDASTNYDTVSEGMGYGLLLAVCFNEQALFDDLYRYVKSHFNGNGLMHWHIDANNNVT
SHDGGDGAATDADEDIALALIFADKLWGSSGAINYGQEARTLINNLYNHCVEHGSYVLKPGDRWGGSSVTNPSYFAPA
WYKVYAQYTGDTRWNQVADKCYQIVEEVKKYNNGTGLVPDWCTASGTPASGQSYDYKYDATRYGWRTAVDYSWFG
DQRAKANCDMLTKFFARDGAKGIVDGYTIQGSKISNNHNASFIGPVAAASMTGYDLNFAKELYRETVAVKDSEYYGYYG
NSLRLLTLLYITGNFPNPLSDLSGQPTPPSNPTPSLPPQVVYGDVNGDGNVNSTDLTMLKRYLLKSVTNINREAADVNRD
GAINSSDMTILKRYLIKSIPHLPY

>SEC_celK (from pSEC_cellulase)

MKLKNTLGLAIGSLVAATSIGAMAMNFRRMLCAAIVLTIVLSIMLPSTVFALEDKSSKLPDYKNDLLYERTFDEGLCFPWH
TCEDSGGKCDFAVVDVPGEPGNKAFRLTVIDKGQNKWSVQMRHRGITLEQGHTYTVRFTIWSDKSCRVYAKIGQMGE
PYTEYWNNNWNPFNLTPGQKLTVEQNFTMNYPTDDTCEFTFHLGGELAAGTPYYVYLDDVSLYDPRFVKPVEYVLPQP
DVRVNQVGYLPFAKKYATVVSSSTSPLKWQLLNSANQVVLEGNTIPKGLDKDSQDYVHWIDFSNFKTEGKGYYFKLPTV
NSDTNYSHPFDISADIYSKMKFDALAFFYHKRSGIPIEMPYAGGEQWTRPAGHIGIEPNKGDTNVPTWPQDDEYAGRP
QKYYTKDVTGGWYDAGDHGKYVVNGGIAVWTLMNMYERAKIRGIANQGAYKDGGMNIPERNNGYPDILDEARWEI
EFFKKMQVTEKEDPSIAGMVHHKIHDFRWTALGMLPHEDPQPRYLRPVSTAATLNFAATLAQSARLWKDYDPTFAAD
CLEKAEIAWQAALKHPDIYAEYTPGSGGPGGGPYNDDYVGDEFYWAACELYVTTGKDEYKNYLMNSPHYLEMPAKM
GENGGANGEDNGLWGCFTWGTTQGLGTITLALVENGLPATDIQKARNNIAKAADRWLENIEEQGYRLPIKQAEDERG
GYPWGSNSFILNQMIVMGYAYDFTGNSKYLDGMQDGMSYLLGRNGLDQSYVTGYGERPLQNPHDRFWTPQTSKKF
PAPPPGIIAGGPNSRFEDPTITAAVKKDTPPQKCYIDHTDSWSTNEITVNWNAPFAWVTAYLDEIDLITPPGGVDPEEPE
VIYGDCNGDGKVNSTDAVALKRYILRSGISINTDNADVNADGRVNSTDLAILKRYILKEIDVLPHK

>TAT_celK (from pTAT_cellulase)

MSRDTGDNLDRNQSGNLPMANVMDAYLSRRSVMRGSLGAAIAMIAGTGLTGCFDGGGSDMNFRRMLCAAIVLTIVL
SIMLPSTVFALEDKSSKLPDYKNDLLYERTFDEGLCFPWHTCEDSGGKCDFAVVDVPGEPGNKAFRLTVIDKGQNKWSV
QMRHRGITLEQGHTYTVRFTIWSDKSCRVYAKIGQMGEPYTEYWNNNWNPFNLTPGQKLTVEQNFTMNYPTDDTCE

FTFHLGGELAAGTPYYVYLDDVSLYDPRFVKPVEYVLPQPDVRVNQVGYLPFAKKYATVVSSSTSPLKWQLLNSANQVV
LEGNTIPKGLDKDSQDYVHWIDFSNFKTEGKGYYFKLPTVNSDTNYSHPFDISADIYSKMKFDALAFFYHKRSGIPIEMPY
AGGEQWTRPAGHIGIEPNKGDTNVPTWPQDDEYAGRPQKYYTKDVTGGWYDAGDHGKYVVNGGIAVWTLMNMY
ERAKIRGIANQGAYKDGGMNIPERNNGYPDILDEARWEIEFFKKMQVTEKEDPSIAGMVHHKIHDFRWTALGMLPHE
DPQPRYLRPVSTAATLNFAATLAQSARLWKDYDPTFAADCLEKAEIAWQAALKHPDIYAEYTPGSGGPGGGPYNDDYV
GDEFYWAACELYVTTGKDEYKNYLMNSPHYLEMPAKMGENGGANGEDNGLWGCFTWGTTQGLGTITLALVENGLP
ATDIQKARNNIAKAADRWLENIEEQGYRLPIKQAEDERGGYPWGSNSFILNQMIVMGYAYDFTGNSKYLDGMQDGM
SYLLGRNGLDQSYVTGYGERPLQNPHDRFWTPQTSKKFPAPPPGIIAGGPNSRFEDPTITAAVKKDTPPQKCYIDHTDS
WSTNEITVNWNAPFAWVTAYLDEIDLITPPGGVDPEEPEVIYGDCNGDGKVNSTDAVALKRYILRSGISINTDNADVNA
DGRVNSTDLAILKRYILKEIDVLPHK

>bglC from *Thermobifida fusca* (from *P. putida* KT2440_P$_{EM7}$-cbh)

MTSQSTTPLGNLEETPKPDIRFPSDFVWGVATASFQIEGSTTADGRGPSIWDTFCATPGKVENGDTGDPACDHYNRYR
DDVALMRELGVGAYRFSIAWPRIQPEGKGTPVEAGLDFYDRLVDCLLEAGIEPWPTLYHWDLPQALEDAGGWPNRDT
AKRFADYAEIVYRRLGDRITNWNTLNEPWCSAFLGYASGVHAPGRQEPAAALAAAHHLMLGHGLAAAVMRDLAGQA
GRSVRIGVAHNQTTVRPYTDSEADRDAARRIDALRNRIFTEPLVKGRYPEDLIEDVAAVTDYSFVQDGDLKTISANLDM
MGVNFYNPSWVSGNRENGGSDRLPDEGYSPSVGSEHVVEVDPGLPVTAMGWPIDPTGLYDTLTRLANDYPGLPLYIT
ENGAAFEDKVVDGAVHDTERIAYLDSHLRAAHAAIEAGVPLKGYFVWSFLDNFEWAWGYSKRFGIVHVDYESQTRTV
KDSGWWYSRVMRNGGIFGQE
